# Genetic screening for mutants with altered seminal root numbers in hexaploid wheat using a high-throughput root phenotyping platform

**DOI:** 10.1101/364018

**Authors:** Oluwaseyi Shorinola, Ryan Kaye, Guy Golan, Zvi Peleg, Stefan Kepinski, Cristobal Uauy

**Author notes:** **Corresponding Authors: Oluwaseyi Shorinola,** Bioscience Eastern and Central Africa - International Livestock Research Institute, Nairobi, P O Box 30709, Kenya. **Cristobal Uauy,** John Innes Centre, Norwich Research Park, Norwich, NR4 7UH, UK.

## Abstract

Roots are the main channel for water and nutrient uptake in plants. Optimisation of root architecture provides a viable strategy to improve nutrient and water uptake efficiency and maintain crop productivity under water-limiting and nutrient-poor conditions. We know little, however, about the genetic control of root development in wheat, a crop supplying 20% of global calorie and protein intake. To improve our understanding of the genetic control of seminal root development in wheat, we conducted a high-throughput screen for variation in seminal root number using an exome-sequenced mutant population derived from the hexaploid wheat cultivar Cadenza. The screen identified seven independent mutants with homozygous and stably altered seminal root number phenotypes. One mutant, Cadenza0900, displays a recessive extra seminal root number phenotype, while six mutants (Cadenza0062, Cadenza0369, Cadenza0393, Cadenza0465, Cadenza0818 and Cadenza1273) show lower seminal root number phenotypes most likely originating from defects in the formation and activation of seminal root primordia. Segregation analysis in F_2_ populations suggest that the phenotype of Cadenza0900 is controlled by multiple loci whereas the Cadenza0062 phenotype fits a 3:1 mutant:wild-type segregation ratio characteristic of dominant single gene action. This work highlights the potential to use the sequenced wheat mutant population as a forward genetic resource to uncover novel variation in agronomic traits, such as seminal root architecture.

## INTRODUCTION

The 1960’s “Green Revolution” demonstrated the impact that changes to plant architecture in major crops like wheat and rice can have on increasing food production (Hedden 2003). While the Green Revolution focused on improving shoot architecture, it did not optimise root architecture, in part because selection was primarily for performance under management regimes involving high rates of fertiliser application (Lynch 2007). In addition to providing anchorage, the root is the main channel for water and nutrient uptake in crops and serves as an interface for symbiotic interaction with the soil microbiome. Roots are often considered as the hidden and neglected other-half of plant architecture and have not been a direct target for selection during early wheat domestication and in modern wheat breeding programmes (Waines and Ehdaie 2007).

In many environments, water availability is the main factor defining crop rotations and performance. Projections on future climate predict more variable weather events relating to the timings and intensity of precipitations which could negatively affect food security (Cattivelli et al. 2008; Rojas et al. 2019). Optimising root system architecture (RSA) for improved nutrient and water uptake under these uncertain scenarios provides a rational approach to help achieve future food and nutrition security.

The wheat root system is comprised of two main root types, seminal (embryonic) and nodal (post-embryonic) roots, that develop at different times. (Manske and Vlek 2002). As the first root type that emerges, seminal roots are entirely responsible for nutrient and water uptake in seedlings. Seminal roots are therefore important for seedling vigour and early plant establishment which also determines competitiveness against weeds. Nodal roots on the other hand are shoot-borne and develop soon after tillering to provide anchorage and support resource uptake especially during the reproductive stage of wheat growth (Manske and Vlek 2002).

Despite their early establishment, seminal roots remain functionally active through to the reproductive stage and may grow up to 2 m in length (Manschadi et al. 2013; Araki and Iijima 2001). They have also been shown to have similar nutrient uptake efficiency as nodal roots in wheat and contribute to yield potential especially under conditions of low soil moisture where nodal roots may not grow (Weaver and Zink 1945; Sebastian et al. 2016). Given their importance, seminal root traits, such as number and angle, have been linked to adaptive responses under water limiting conditions (Manschadi et al. 2008; Cane et al. 2014; Golan et al. 2018). Steep seminal root angle has been associated with increased soil water exploration at depth which is beneficial in drought condition where topsoil moisture is depleted (Richard et al. 2015; Olivares-Villegas et al. 2007; Manschadi et al. 2008).

Seminal roots develop from the root primordia in the embryo of a germinating wheat seed. There is genetic variation among wheat genotypes for the number of seminal roots that develop, this can range from three to six seminal roots per plant dependent on cultivar (Araki and Iijima 2001; Robertson et al. 1979; Golan et al. 2018). Typically, the seminal root system consists of a primary root that emerges first followed by two pairs of secondary seminal roots that emerge sequentially. A sixth seminal root may also develop in some wheat varieties. A few quantitative trait loci (QTL) have been identified to underlie variation in seminal root number in wheat germplasm (Atkinson et al. 2015; Maccaferri et al. 2016; Ren et al. 2012; Sanguineti et al. 2007; Ma et al. 2017; Iannucci et al. 2017). However, many of these QTLs are defined to broad intervals which makes their genetic dissection difficult and their use in breeding limited. Unlike other cereals (e.g. rice, maize), only one gene controlling root system architecture (RSA), *VERNALIZATION1* (*VRN1*; Voss-Fels et al. 2018), has been identified in wheat.

This delay in identifying genetic loci controlling root traits is most likely due to a series of factors which makes genetic analyses in wheat difficult. Bread wheat is a hexaploid plant with a relatively large (16 Mb) and repeat-rich (>85%) genome comprised of three homoeologous sub-genomes (B, A and D). High sequence similarity in the coding regions of these sub-genomes results in high levels of genetic redundancy that mask the phenotypic effects of underlying natural variation for many traits, including RSA traits (Uauy et al. 2017; Borrill et al. 2015). Also, the “out-of-sight” nature and extreme phenotypic plasticity of roots under native field conditions makes root phenotyping difficult, cumbersome and time-consuming (Atkinson et al. 2019).

The use of induced variation has proven useful to uncover novel phenotypes and dissect genetic pathways underlying complex phenotypes in plants (Parry et al. 2009). Our current understanding of the genetic determinants regulating root development in many cereals have almost entirely stemmed from the isolation and characterisation of mutants defective in one or more RSA traits (Reviewed in Coudert et al. 2010; Hochholdinger et al. 2018; Marcon et al. 2013). Despite this potential, the use of mutant populations to study the genetic control of root development in wheat has not hitherto been exploited. The recent development of an *in-silico* platform for the rapid identification of mutations in 1,200 ethyl methanesulfonate (EMS) mutagenized lines in the UK hexaploid wheat cultivar ‘Cadenza’ now makes large-scale reverse and forward genetic investigation of traits more feasible in wheat (Krasileva et al. 2017). Progress has also been made on the root phenomics front, with the development of fast, low-cost, and flexible two-dimensional (2D) root phenotyping pipelines with sufficient throughput for phenotyping large populations (Selvara et al. 2013; Atkinson et al. 2019; Adeleke et al. 2019).

Taking advantage of these new developments, we implemented a relatively high-throughput root phenotyping pipeline to conduct a forward genetic screen for variation in seminal root number using a subset of the exome-sequenced Cadenza mutant population. From this work, we describe the identification and characterisation of novel hexaploid bread wheat mutants with decreased and increased numbers of seminal roots.

## MATERIALS AND METHODS

### Germplasm

#### Mutant population for primary screens

A hexaploid wheat mutant population was previously developed by EMS treatment of the UK bread wheat cultivar Cadenza (Krasileva et al. 2017; Rakszegi et al. 2010). In this study, we used 645 exome-sequenced mutants from this population (Krasileva et al. 2017). These mutant lines were selected based on the criteria that they show greater than 90% germination rate during a seed multiplication that was conducted in the field. To obtain homogenous phenotypes and reduce the variation from segregating mutations, single spikes harvested from field-grown M_4_ plants were individually threshed and derived M_5_ seeds (40 - 50 seeds) from each spike were divided for use in forward genetic Screen A and Screen B described below (Figure 1).

**Figure 1:**
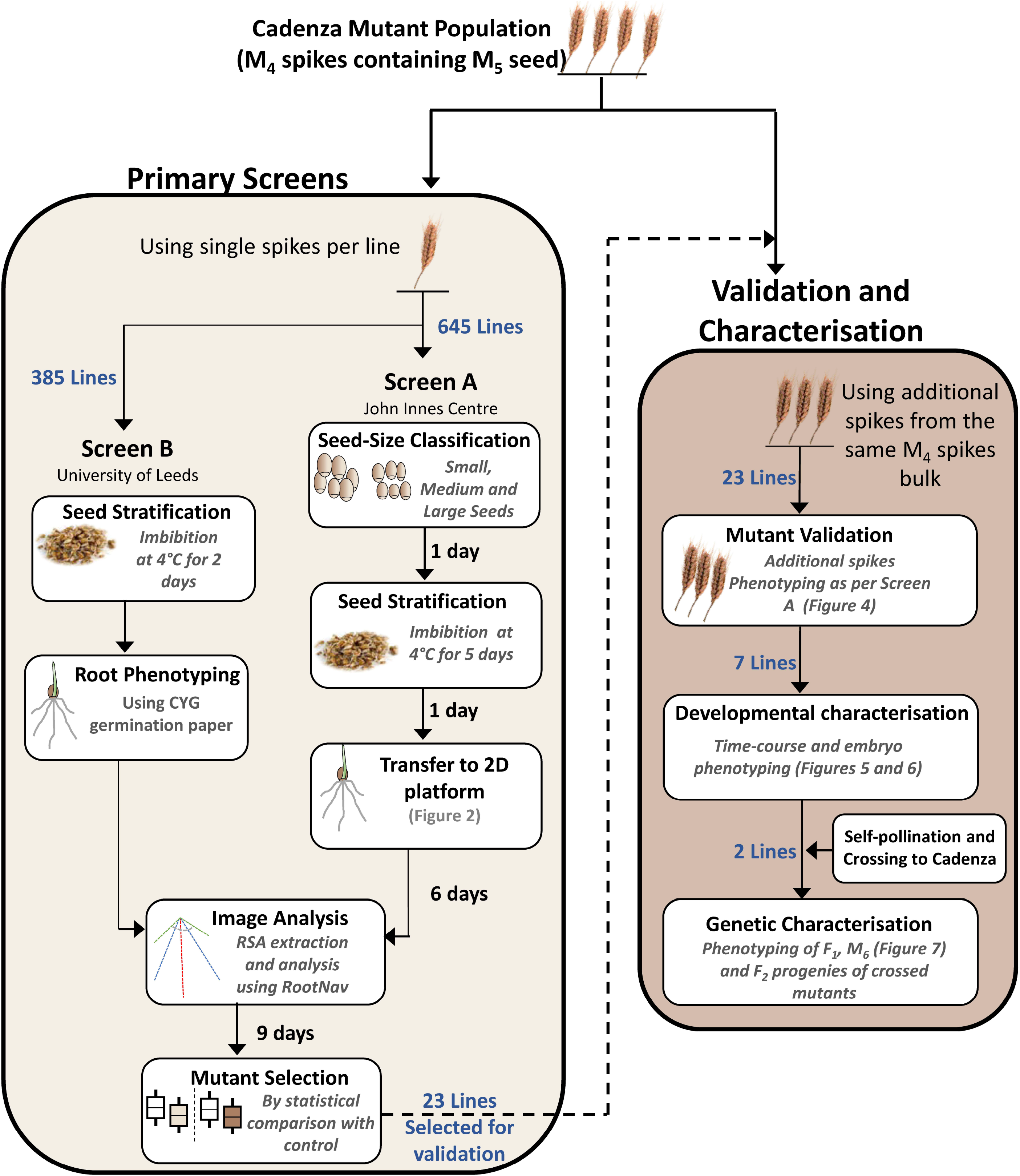
Schematic workflow of the forward genetic screens, and validation and characterisation experiments. Single spikes for each mutant were first phenotyped in the primary screens (light brown box) and selected mutants were subsequently phenotyped in the validation and characterisation experiments (dark brown box). The number of lines used in each experiment are indicated in blue text. Times indicated in Screen A are an approximation of the time taken to set-up the high-throughput Screen A per phenotyping batch (~1,800 plants) in 8-hours work days.

#### Germplasm for mutant validation and characterisation

Three additional M_4_ spikes (containing M_5_ seed) from the same field grown samples as those used for the primary screens (Figure 1) were used to further validate the phenotypes of the seven mutants identified in the primary screens (described in results). The spikes were individually threshed and M_5_ seeds from each spike were phenotyped separately. Two validated mutants (Cadenza0900 and Cadenza0062) were further selected for genetic characterisation: M_5_ plants were grown to maturity for cross-pollination with wild-type Cadenza to generate F_1_ hybrids which were subsequently self-pollinated to generate F_2_ progenies. M_5_ plants of the selected mutants were also self-pollinated to generate M_6_ seeds to characterise the stability of their phenotypes in the subsequent generation.

### High-throughput Seminal Root Phenotyping

Two similar and independent screens (Screen A and B) were conducted concurrently on subsets of the Cadenza mutant population to identify lines with altered seminal root numbers (Figure 1). All 645 lines were phenotyped at high-throughput in Screen A, while only the first 385 of the 645 lines (in numerical order) were phenotyped in Screen B due to more limited throughput. For mutant lines phenotyped in both screens, M_5_ seeds from the same field-grown M_4_ spike were used in both screens as described in the germplasm section above.

#### Screen A

This screen was carried out at the John Innes Centre, UK, using a custom 2D root phenotyping platform based on the protocol described by Atkinson et al. (2015) with some modifications to increase the throughput from 360 to 1,800 seedlings per run. This screen also took seed size effect on RSA traits into consideration. For each mutant line, M_5_ seeds from a M4 single spike were first size-stratified into large, medium or small seed by passing the seeds through two sets of calibrated graduated sieves with 2.8 mm and 3.35 mm mesh sizes. Large-sized seeds were collected above the 3.35 mm sieve, medium-sized seeds collected between the 2.8 mm and 3.35 mm sieves, and small-sized seeds were collected below the 2.8 mm sieve. Each mutant line was thus designated as either being Large (524 lines), Medium (119 lines) or Small (2 lines) based on the size group where most of its seeds were defined. Only seeds representative of each mutant line size classification were used. The two mutants with small seed size were phenotyped for variation in seminal root number compared to wild-type Cadenza, but were not included in the analysis of seed size effect on RSA due to the small sample size. Seeds (15 per mutant line) were surface sterilized by rinsing in 5% (v/v) Sodium Hypochlorite (Sigma Aldrich, UK) for 10 mins and were rinsed with water three times before being imbibed in 1.75 mL of water for 5 days at 4°C to ensure uniform germination. Of these, 10 seeds per mutant were placed crease facing down into individual growth pouches made from a sheet of germination paper (21.5 cm x 28 cm; Anchor Paper Company, St Paul, MN, USA) clipped to a black polythene sheet (22 cm x 28 cm, 75 μm thickness, Cransford Polythene LTD, Suffolk, UK) using an acrylic rod and 18 mm fold clip (Figure 2A). The growth pouches were suspended in an upright position in plastic storage boxes (120 cm x 27 cm x 36 cm, Really Useful Product, West Yorkshire, UK) with 60 pouches per box (Figure 2B). The sides of the box were covered in black plastic sticky back cover film to block out light from the roots of the developing seedlings. Each box was filled with 10 L of half-strength Hoaglands growth solution containing (Hoagland and Arnon 1950): NH_4_H_2_PO_4_, 0.6 g; Ca(NO_3_)_2_, 3.3 g; MgSO_4_, 1.2 g; KNO_3_, 1.0 g; H_3_BO_3_, 14.3 mg; Cu_2_SO_4_, 0.4 mg; MnCl_2_(H_2_O)_4_, 9.1 mg; MoO_3_, 0.1 mg; ZnSO_4_, 1.1 mg; KHCO_3_, 2.0 g, Ferric Tartrate, 2.8 g. The base of each pouch was suspended in the growth solution to supply nutrients to the developing seedling through capillary action. A randomised complete block design was adopted with each line replicated across 10 different boxes (blocks). The phenotyping boxes were placed in a controlled environment room under long day conditions with 16h light (250–400 mmol) at 20°C, 8h darkness at 15°C and at 70% relative humidity (Figure 2C). After 7 days of growth, pouches were taken out of the phenotyping box; placed on a copy stand and the black plastic sheet covering the germination paper was gently pulled back to reveal the roots. Images of the roots were taken with a Nikon D3400 DSLR Camera fitted to the copy stand (Figure 2D). Phenotyping of the mutant population was done over 12 experiments with 60 lines (59 mutants and a Cadenza control) phenotyped per experiment. Mutants of the same seed-size group were phenotyped together – large sized mutants in experiment 1 - 9 and small/medium sized mutants in experiment 10 - 12. Cadenza seeds representative of the seed-size group for each experiment were used as controls. The 12 experiments were completed across five rounds of phenotyping with 2 - 3 experiments (120 – 180 lines) set up per round. For 96% of the mutants examined, we successfully imaged the roots of 8 - 10 plants. In the remain 4% of the mutants, only 3 - 7 plants could be images due to poor seed germination or poor seedling growth on the pouch. In total, 6,240 (6,127 mutant and 113 Cadenza) plants were phenotyped. The same phenotyping set-up was used for the M_5_ validation experiments and to characterise M_6_, F_1_ and F_2_ progenies of the two mutants selected for genetic characterisation.

**Figure 2:**
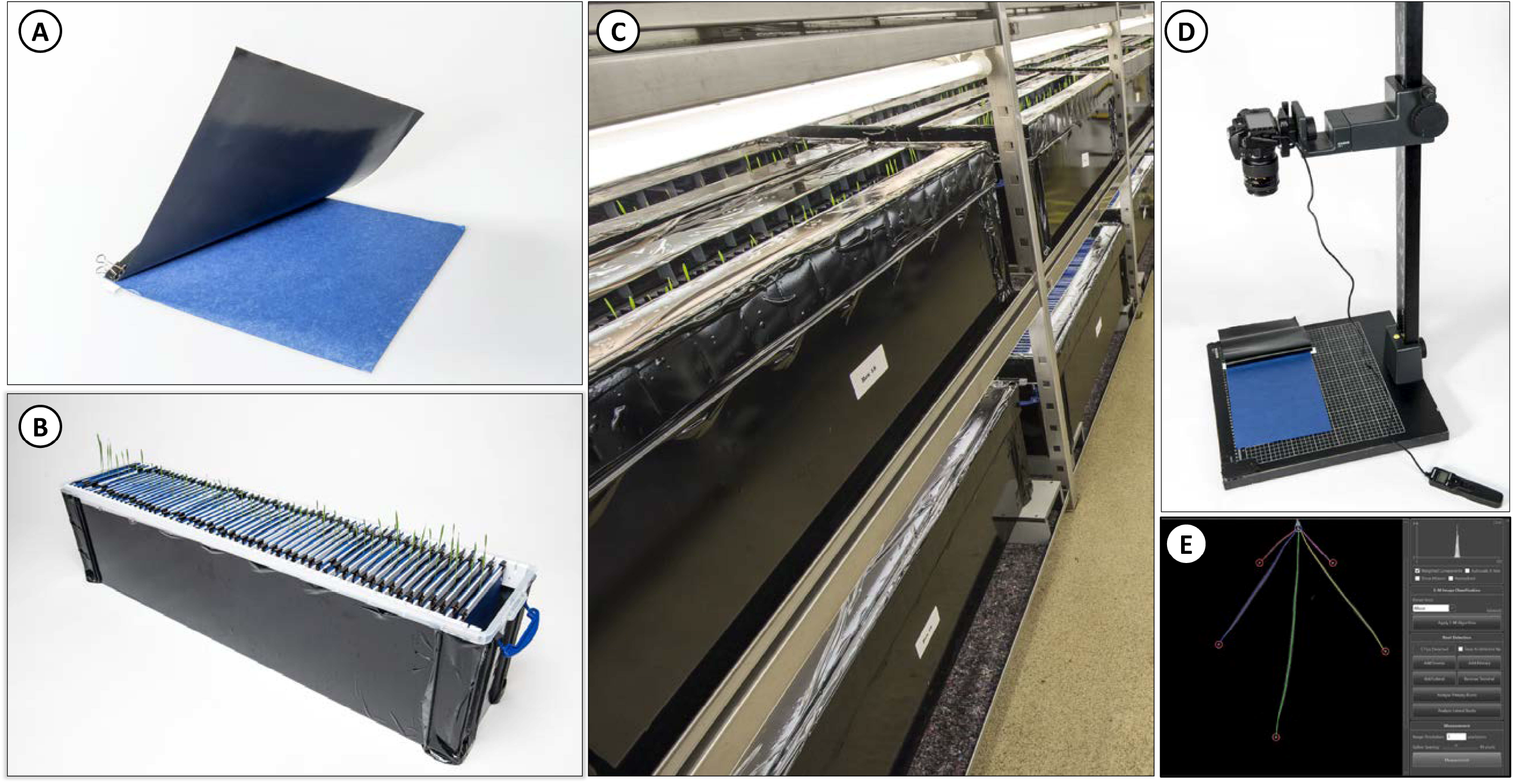
Root phenotyping set-up used for screen A: (A) Growth pouch showing blue blotter germination paper and cover plastic sheet. (B) Phenotyping box containing growth pouch (60 pouches per box) and nutrient solution at the bottom. (C) Root phenotyping in controlled environment room. (D) Nikon D3400 DSLR Camera mounted on copy stand for root imaging. (E) Digital extraction of root architecture using RootNav.

#### Screen B

A second screen was conducted in parallel at the University of Leeds, UK, using commercial CYG paper pouches (Mega-International, Minnesota, USA). Due to the more limited throughput of this screen, only the first 385 of the 645 mutant lines (in numerical order) were examined in this screen. For each mutant line, 10 visually uniform M_5_ seeds were selected and placed onto moist filter paper in a 90 mm round petri dish. Petri dishes were wrapped in aluminium foil to exclude light and placed at 4°C for 2 days, seeds were placed crease side down into individual CYG seed germination pouches with the bottom removed to allow wicking of growth solution from a reservoir of media. Pouches were wrapped in aluminium foil in batches of five to exclude light from the roots and were placed upright in a reservoir of full strength Hoaglands No 2 growth solution (NH_4_H_2_PO_4_, 115.03 mg; Ca(NO_3_)_2_, 656.4 mg; MgSO_4_, 240.76 mg; KNO_3_, 606.6 mg; H_3_BO_3_, 2.86 mg; Cu_2_SO_4_, 0.08 mg; MnCl_2_(H_2_O)_4_, 1.81 mg; MoO_3_, 0.016 mg; ZnSO_4_, 0.22 mg; Ferric Tartrate, 5 mg. per Litre). Pouches were placed in long day conditions (as in screen A) and were grown for 5 days before roots were imaged. For imaging, pouches were placed onto a copy stand; the front of the pouch carefully removed, and the root system imaged using a Sony Cybershot DSC-RX100. Plants were screened in rounds of twenty lines with a total of 200 plants per round.

### Image Analysis

High-resolution images captured from the phenotyping were pre-processed (rotated, cropped and compressed) using ImageJ (https://imagej.nih.gov/index.html) and Caesium image compressor before being processed in RootNav (Pound et al. 2013). Captured root architectures were imported into a RootNav viewer database for measurement of RSA traits using standard RootNav functions (Figure 2E).

### Anatomical Characterisation of Seminal Root Primordia

Embryos of mutants validated to have reduced seminal root number phenotypes were examined using the method described by Golan et al. (2018). In brief: embryos from mature dry grains were fixed in FAA solution (10% formaldehyde, 5% acetic acid, 50% ethanol, and 35% distilled water by volume) overnight and dehydrated at room temperature in a graded ethanol series (30 min each, in 50%, 70%,90%, 95%, and 100% ethanol). Then, embryos were cleared in xylene, embedded in paraffin and sectioned (5 μM) using a microtome (Leica Biosystems, Germany). Cross sections were de-paraffinized with histoclear, rehydrated and stained with Harris Hematoxylin. A stereo microscope (SZX16, Olympus, Tokyo, Japan) was used for imaging.

### Statistical Analysis

All statistical analyses were performed in R 3.5.1 (R Core Team 2018) and Minitab 17 statistical software. Statistically significant seminal root number difference in the primary screening experiments (Screen A and B) was determined by ANOVA using a Dunnett’s comparison within each phenotyping experiment with the Cadenza plants in each experiment used as control. Adjusted probability values of *P* < 0.05 were considered statistically significant. Statistically significant root architectural difference in the validation experiment as well as in the M_6_ and F_1_ phenotyping experiments were based on Student’s t-test comparison of individual spike/line to the Cadenza control. A Chi-square test was used to examine the goodness of fit of the segregation pattern observed in the F_2_ progenies to patterns expected for single recessive or dominant gene action.

### Data Availability

Seeds of mutant lines reported in this study can be ordered through the SeedStor site at www.seedstor.ac.uk. Supplemental materials (tables and figures) are available at *FigShare*. Table S1 contains information on mutants with significantly different seminal root number phenotypes to Cadenza from Screen A. Table S2 contains information on all the mutants phenotyped in both Screen A and B. Figure S1 shows the relationship between seminal root number and total root length in Screen A. Figure S2 and Figure S3 show representative images and embryo size measurement of mutants with validated altered seminal root number phenotypes, respectively. All the root images from Screen A (6,240 images) including the original RootNav measurements for different root traits are publicly available on Zenodo (https://doi:10.5281/zenodo.3270726) for download and reuse.

## RESULTS

### Identifying induced variation for seminal root number in hexaploid wheat

We implemented a 2D root phenotyping pipeline suitable for large-scale phenotyping at a throughput of 1,800 seedlings per run (Figure 1 and 2). Using this platform, we performed a forward genetic screen for variation in seminal root number using 645 seed-size stratified (small, medium and large; see Methods) M_5_ mutants from the exome-sequenced Cadenza mutant population (Krasileva et al. 2017). In our screen, Cadenza mainly displayed five seminal roots (4.9 ± 0.05) including a primary seminal root (SR_1_), as well as first (SR_2,3_) and second (SR_4,5_) pairs of seminal roots (Figure 3A). We observed variation in seminal root number in the mutant population, with seminal root number ranging from 1 to 7 in individual plants and mean seminal root number per mutant (with *n* ≥ 4 plants per mutant) ranging from 2.9 to 5.9.

**Figure 3:**
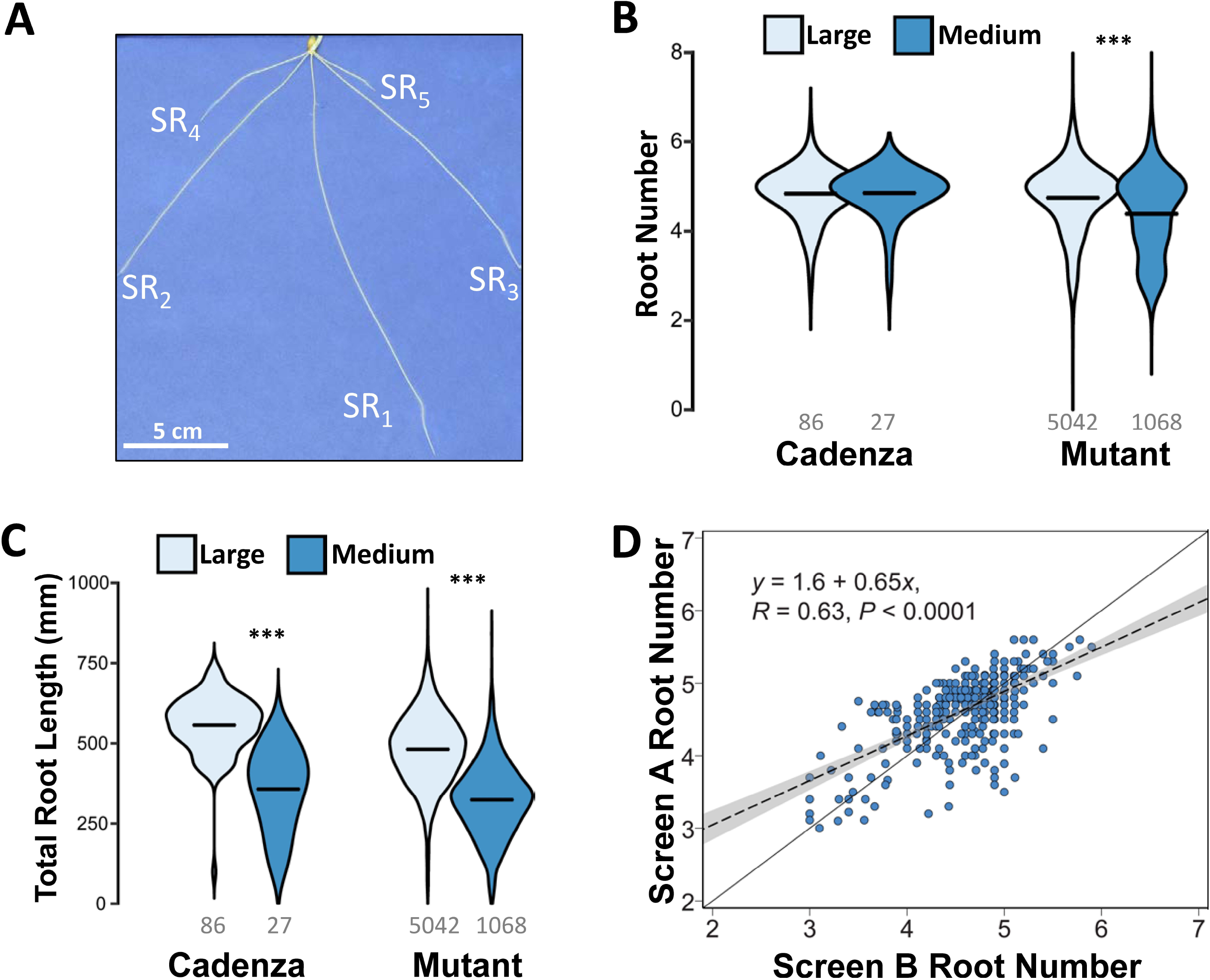
Variation in seminal root number in the Cadenza mutant population. (A) Seminal root architecture of a Cadenza seedling showing the primary root (SR_1_) and the first (SR_2,3_) and second pairs of seminal roots (SR_4,5_). (B-C) Distribution of the seminal root number (B) and total root length measurements (C) phenotypes observed in Screen A across large and medium seed-size groups. The mid-line represents the mean of the distribution. The number of plants in each seed-size group are indicated below each distribution. *** indicates *P* <0.0001 for statistical comparisons between the seed-size groups. (D) Regression of the seminal root number phenotypes observed in primary Screens A and B using a linear model. Only lines phenotyped in both screens are shown. The regression line of the two screens (dotted diagonal line) is compared to a hypothetical perfect correlation (solid line) between the screens.

Within the mutant lines, seed size groups (large and medium) showed significant difference in seminal root number (*P* < 0.0001) and total root length (*P* < 0.0001). Mutants with large and medium-sized grains had an average seminal root number of 4.7 and 4.4, respectively, and total root length of 481 mm and 324 mm, respectively (Figure 3B-C). In the Cadenza wild-type, significant differences between seed size groups were only observed for total root length. There were only two mutant lines with small seed size and these were not included in the analysis of seed size effect on seminal root traits. Across all mutant lines, we observed a significant positive correlation (*R* = 0.56, *P* <0.0001) between the number of seminal roots and the total root length in the population (Figure S1).

Parallel to this, we phenotyped a subset of these Cadenza mutants for similar root traits in an independent screen (Screen B; Figure 1). Given the lower throughput of this approach (See Screen 2 in Methods), only 385 out of the 645 mutant lines were screened. We observed a significant positive correlation between seminal root number measurements in Screen A and Screen B (*R* = 0.63; *P* <0.0001; Figure 3D). The heritability estimate of the seminal root measurement across the two screens was 0.77 suggesting a strong heritable genetic effect in the determination of seminal root number in the Cadenza mutant population.

We first assessed the statistical significance from the mutants studied in Screen A. Dunnett’s multiple comparison identified 52 mutants (8 % of mutants in Screen A) with significantly different number of seminal roots relative to the Cadenza control across the three seed size groups (Table S1). Five of these mutants had significantly higher number of seminal roots with mean seminal root number ranging from 5.7 to 5.9 and modal seminal root number of 6 per mutant (Table S1). The higher seminal root phenotype is mainly driven by the development of an extra root, hereafter referred to as SR_6_. The remaining 47 mutants showed significantly lower number of seminal roots with mean seminal root number per mutant of 2.9 to 4.1 and modal seminal root number between 3 and 5.

Thirty three out of the 52 significant mutants identified in Screen A were also phenotyped as part of Screen B. This included four of the five higher root number mutants and 29 lower root number mutants. Details of individual mutant lines phenotyped in both screens are presented in Table S2. In Screen B we confirmed the significant phenotype of three of the four higher root number mutants in common with Screen A; these lines displayed a mean seminal root number of 5.1 to 5.5 in Screen B. In the case of the 29 lower root number mutants which were investigated in both screens, we confirmed 20 mutants which displayed fewer numbers of seminal roots (less than 4 roots) than Cadenza. Based on the results from the two screens, we selected the three higher and 20 lower root number mutants with consistent phenotypes in both screens for further phenotypic evaluation.

### Altered root number mutants show stable homozygous seminal root number phenotypes

To validate the 23 selected mutants, we phenotyped M_5_ seeds from three additional M_4_ spikes (containing M_5_ seeds) from the same field-grown bulk as the spike used in the primary screen (Figure 1). These M4 spikes originate from successive bulking of multiple M_3_ and M4 plants. Selecting three separate spikes increases the probability of phenotyping plants with independent background mutations thereby providing robust biological replications to examine the stability of the mutations effects and segregation patterns (homozygous or heterozygous).

For seven of the selected mutants including one higher root number (Cadenza0900) and six lower root number mutants (Cadenza0062, Cadenza0369, Cadenza0393, Cadenza0465, Cadenza0818, and Cadenza1273), we observed the altered seminal root number phenotype in the three additional spikes phenotyped (Figure 4, Figure S2). This suggests that the phenotypes of these mutants are consistent across sibling lines and controlled by mutations that were most likely homozygous in the original single M_2_ plant from which the population was derived. For the rest of the 16 mutants, we did not consistently observe the altered seminal root number phenotypes in all three additional spikes. These might represent lines with segregating phenotypes arising from heterozygous mutations in the initial M_2_ plants or false-positive selection in the primary screens; these lines were not studied further.

**Figure 4:**
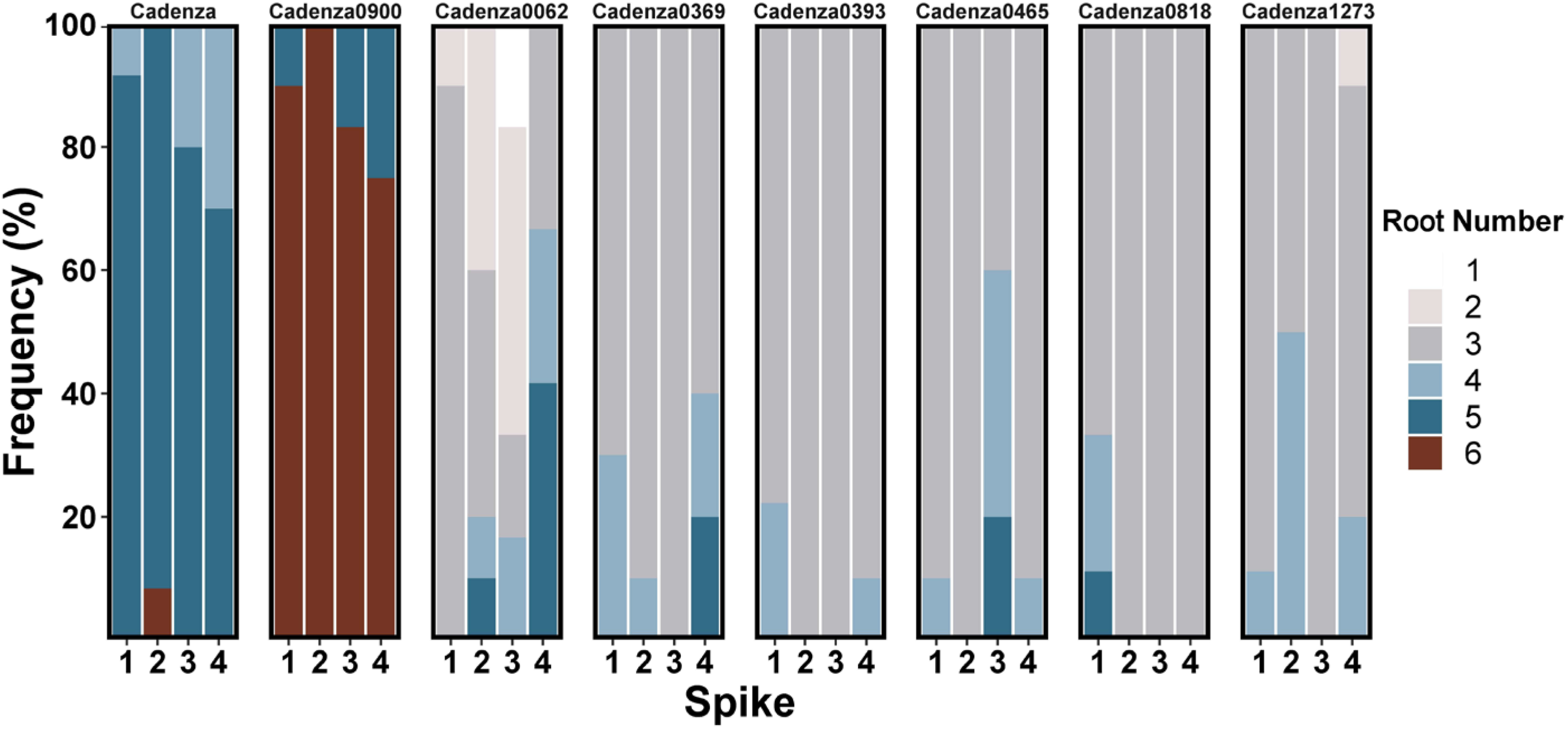
Validated mutants show homozygous seminal root number phenotypes. Seminal root number distribution in wild-type Cadenza and mutants with validated altered seminal root number phenotypes across four spikes phenotyped in primary Screen A (spike 1) and the validation experiments (spikes 2 to 4). The number of plants phenotyped from each spike ranged from four to ten.

We further characterized the seven validated mutants from 1 to 7 days post germination (dpg) to examine when the phenotype was first visible and to identify the seminal root type (primary, first pair or second pair) defective in these mutants (Figure 5). Cadenza showed fully emerged primary root (SR_1_) at 1 dpg, while the first (SR_2,3_) and second pairs (SR_4,5_) of seminal roots emerged at 3 and 5 dpg, respectively. Similar to Cadenza, Cadenza0369, Cadenza0393, Cadenza0465, Cadenza0818, and Cadenza1273 with lower root counts also developed the primary and first pair of seminal roots at 1 dpg and 3 dpg, respectively, but are defective in the development of the second pair of seminal roots (SR_4,5_ or only SR_5_). Contrary to this, Cadenza0900 with higher root count showed a faster rate of seminal root development relative to Cadenza with the primary, first and second pair of seminal roots having emerged by 3 dpg and an extra sixth root emerged at 7 dpg. Cadenza0062 showed a strong dormancy phenotype and was not included in this experiment.

**Figure 5:**
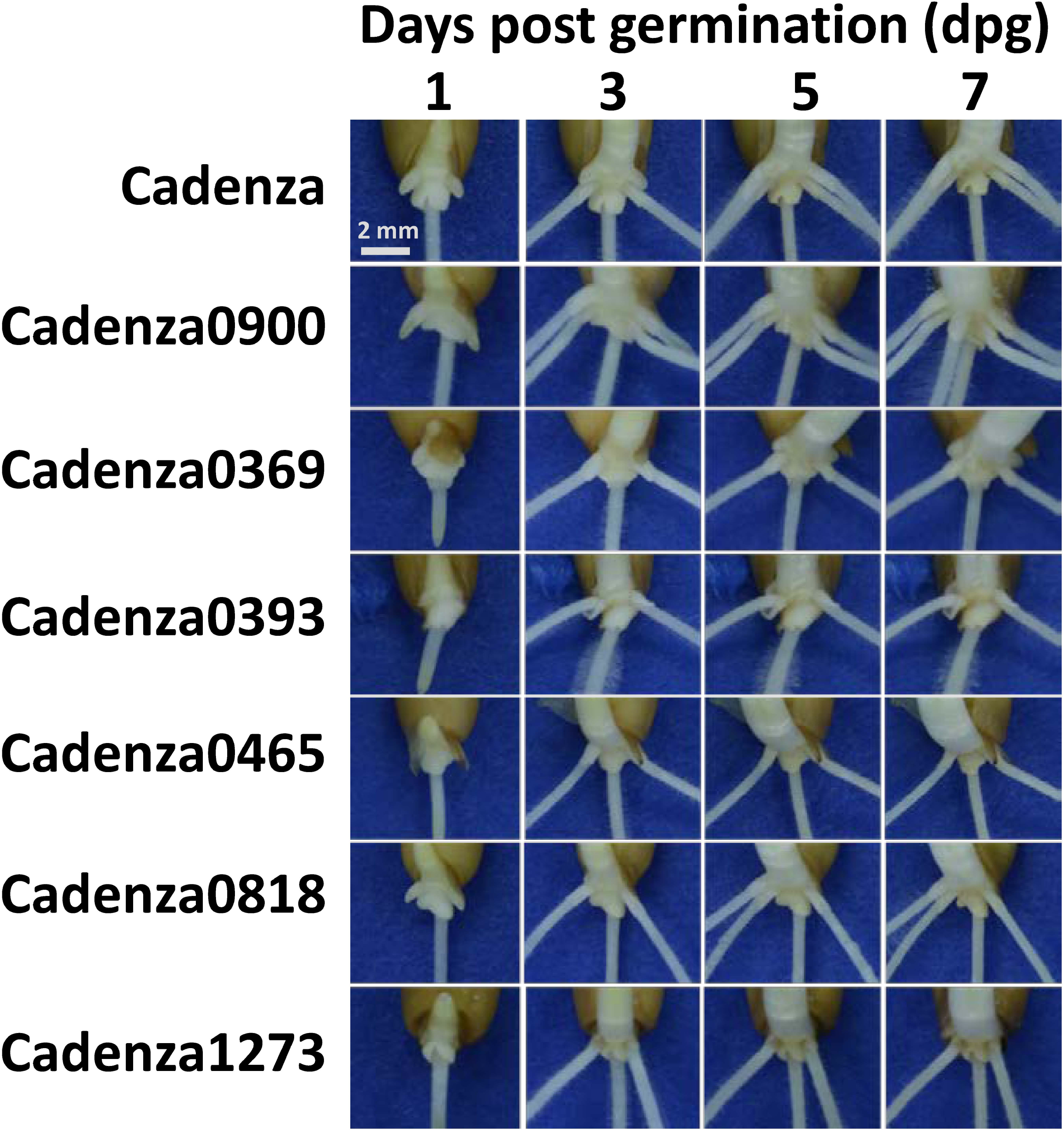
Characterisation of root development in validated mutants. Temporal characterization of seminal root development in the validated mutants at 1, 3, 5 and 7 days post germination (dpg) shows altered SR_4,5_ and SR_6_ seminal root type phenotypes.

### Embryo imaging of the lower root number mutants points to defects in primordia development and growth

Seminal roots emerge from the activation of root primordia that form in the embryo of a developing seed. The primary (SR_1_) and secondary (SR_2,3_ and SR_4,5_) seminal roots emerge from primordia formed in the central portion and sides of the embryo, respectively. Although uncommon, the sixth seminal roots (SR_6_ like in Cadenza0900) are known to emerge from a primordium located in the midpoint between the primordia of the second pair of secondary seminal root (Hoshikawa 1964). Importantly, differential activation of root primordia of SR_4,5_ has been shown to account for lower numbers of seminal roots in some wild wheat species (Golan et al., 2018). We therefore examined if the seminal root phenotypes of the lower root number mutants (Cadenza0062, Cadenza0369, Cadenza0393, Cadenza0465, Cadenza0818, and Cadenza1273) lacking either SR_5_ or SR_4,5_ originate from defects in primordia development and/or https://www.ncbi.nlm.nih.gov/pmc/articles/PMC4169152/ the failure of developed root primordia to activate to become seminal roots. Cadenza (WT) consistently develops five fully formed root primordia (Figure 6). All the lower root number mutants examined showed altered root primordia development compared to Cadenza, with SR_4,5_ (or only SR_5_) primordia either absent or reduced in size (Figure 6). In addition, primordia activity was altered in the mutants, as all mutants had greater number of primordia compared to the number of roots observed in the M_5_ seedlings (Figure 5, Table 1). These measurements provide an initial indication that the defects in the lower root count mutants most likely originate from a combination of both lower number and activity of root primordia in these lines. Embryo size was also significantly lower in four of the six mutants (Cadenza0393, Cadenza0465, Cadenza0818, and Cadenza1273; Figure S3), but it is not clear from these results if the smaller embryo size of these mutants contributes to their lower seminal roots number phenotypes.

**Figure 6:**
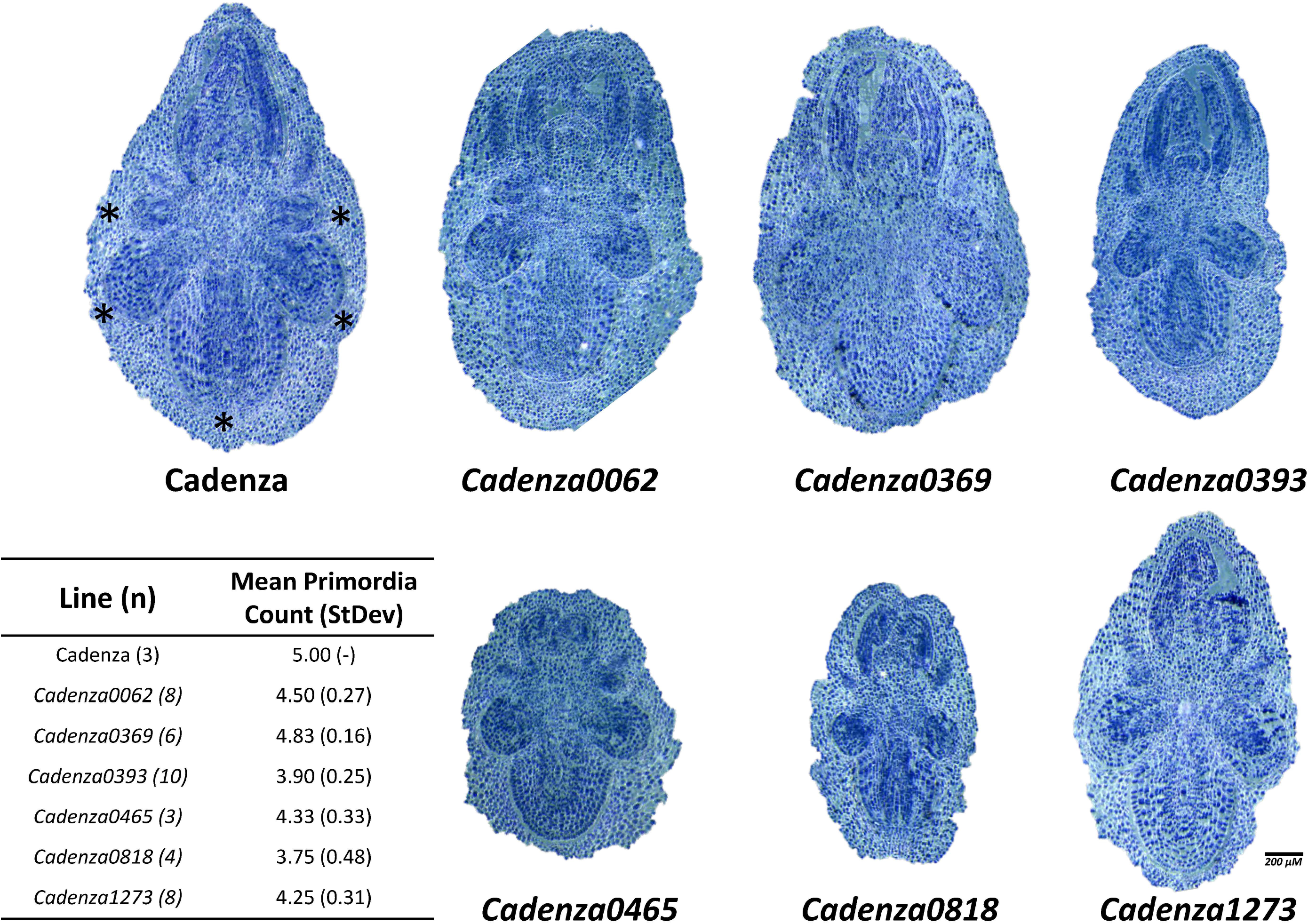
Primordia imaging in lower root number mutants. Longitudinal cross sections of mutants with validated lower seminal root number phenotype prior to seed imbibition. The seminal root primordia of Cadenza are marked by asterisks. The inset table shows the primordia count observed in Cadenza and the lower root number mutants. The number of embryos images are indicated in parenthesis beside each line.

**Table 1:**
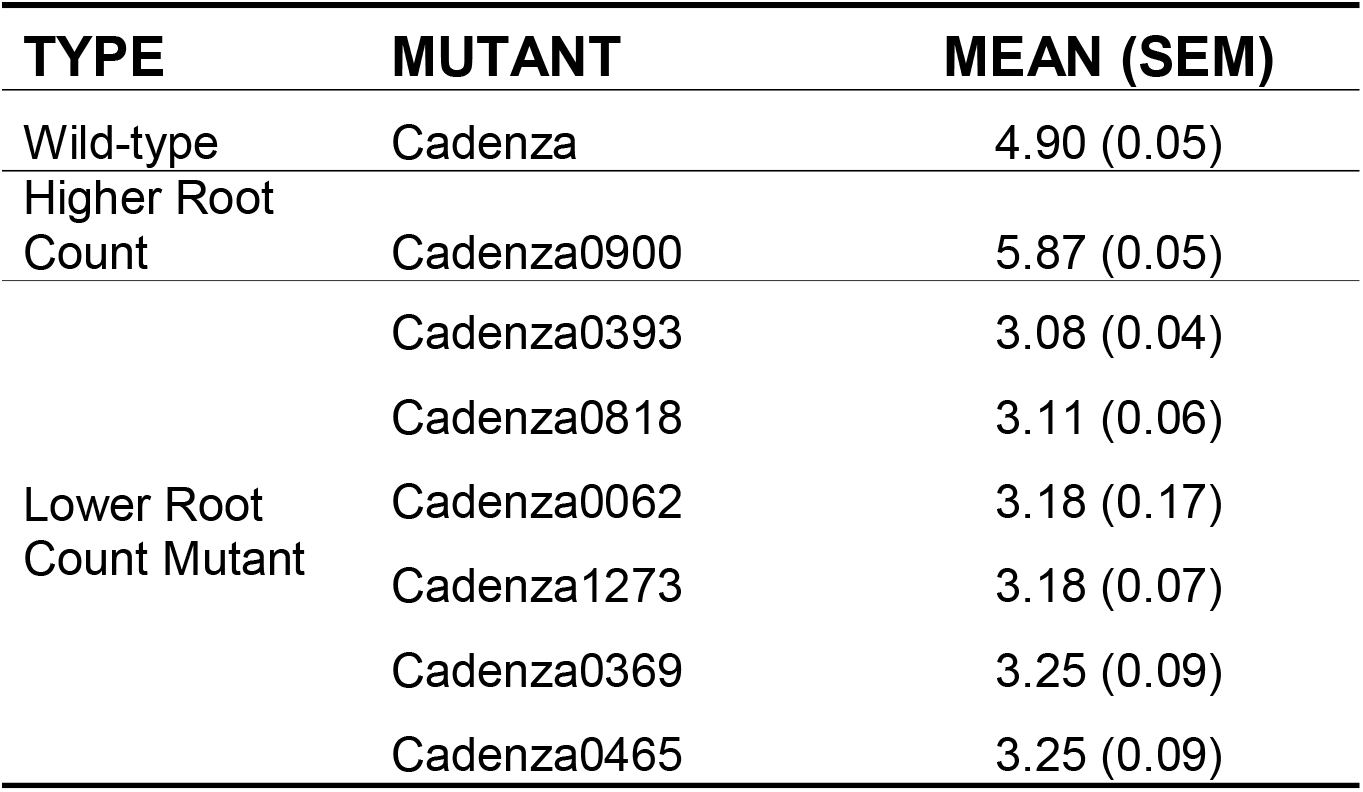
Phenotypic summary of validated altered root number mutants with information on the mutation type

### Genetic characterisation of the altered root number phenotypes

To understand the transgenerational stability and mode of inheritance of the altered root number mutant phenotype, we characterized M_6_ plants of the higher root number mutant (Cadenza0900) and one lower root number mutant (Cadenza0062), as well as F_1_ hybrids derived from crosses of these mutants to Cadenza. M_6_ progenies of Cadenza0900 showed significantly (*P* <0.0001) increased number of seminal roots compared to Cadenza and no significant difference to the phenotypes of it M_5_ parents (Figure 7) with an average root number of 5.73 and more than 73% of the plants having six roots. F_1_ hybrids of Cadenza0900 x Cadenza all had five seminal roots like Cadenza (*P* = 0.19), but significantly lower than their M_5_ plants (*P* < 0.0001; Figure 7). This suggests that the Cadenza0900 phenotype originates from a recessive mutation or is caused by a combination of loci segregating independently.

**Figure 7:**
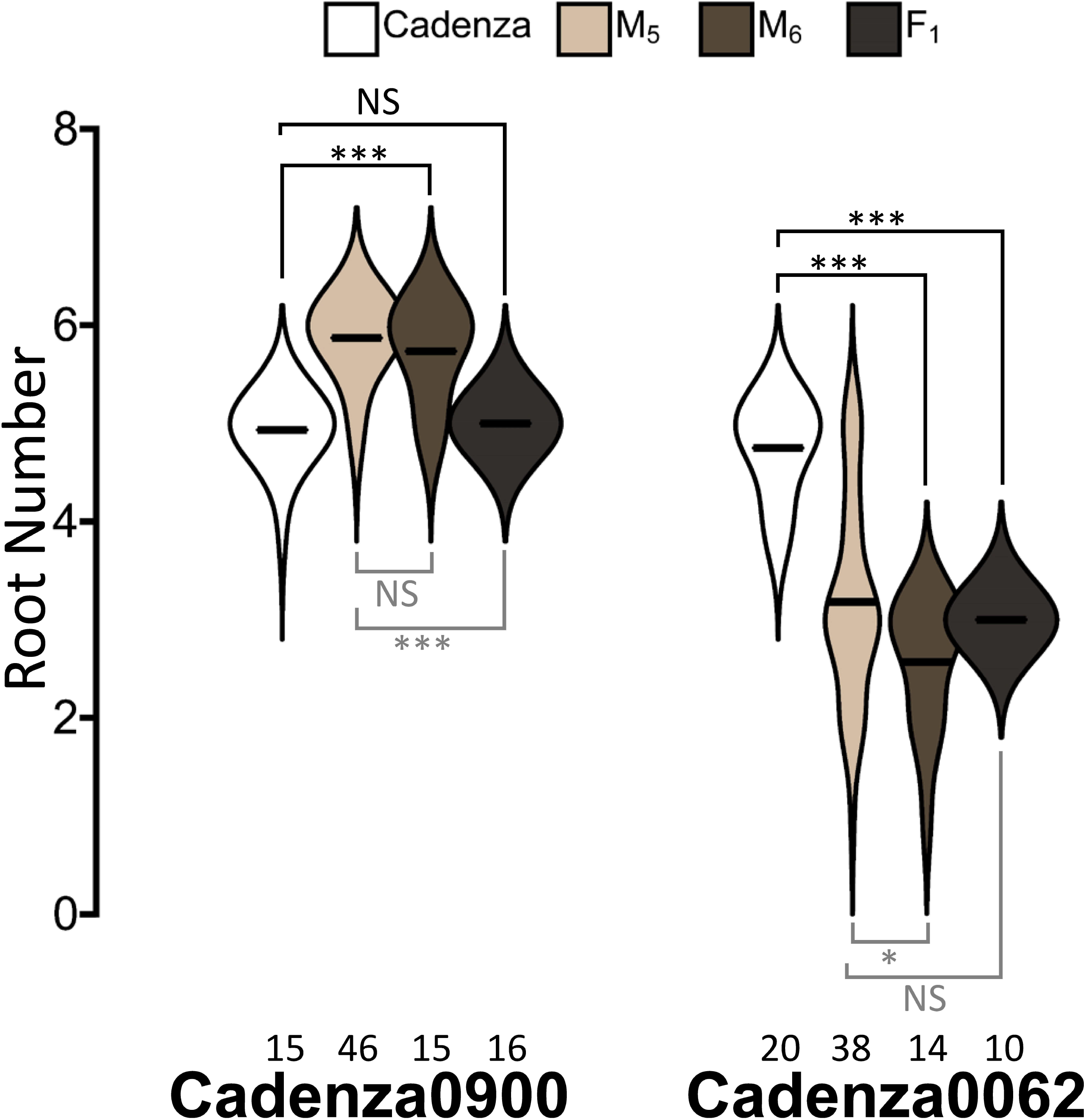
Genetic characterisation of the Cadenza0900 and Cadenza0062 seminal root phenotypes. Seminal root number distribution of M_5_, M_6_ and F_1_ progenies of Cadenza0900 and Cadenza0062 mutants are presented alongside the Cadenza control. The number of plants phenotyped for each genotype is indicated below the distribution. * and *** indicates *P* ≤0.05 and *P* <0.0001, respectively, and the comparison groups are indicated by solid lines; black comparisons to Cadenza and grey comparisons to M_5_ plants. NS indicates non-significant difference.

The M_6_ plants of Cadenza0062 showed a significantly lower number of seminal roots (*P* <0.0001) compared to Cadenza with a mean root number of 2.6 (Figure 7). The seminal root numbers of the M_6_ progenies of Cadenza0062 were also slightly but significantly (*P* = 0.04) lower than the M_5_ plant, most probably due to smaller variation in the seminal root number phenotype in the M_6_ lines. With an average root number of 3.0 (Figure 7), the F_1_ hybrid of Cadenza0062 x Cadenza showed a significantly (*P* <0.0001) lower number of seminal roots compared to Cadenza but a non-significant difference to the original M_5_ plants (Figure 7). This suggests that the Cadenza0062 phenotype is caused by a dominant mutation(s). Unlike the M_5_ plants, the Cadenza0062 x Cadenza F_1_ did not show any reduced germination, suggesting that the dormancy phenotype of Cadenza0062 segregates independently of its altered root number.

To gain further insight into the genetic architecture underlying higher and lower root count phenotypes, we further characterized F_2_ progenies of the Cadenza0900 and Cadenza0062 crosses to Cadenza. We used the chi-square test-statistic to test the goodness of fit of the inheritance pattern of Cadenza0900 and Cadenza0062 phenotypes to those consistent with segregation of single recessive and single dominant traits, respectively. The phenotype of Cadenza0900 F_2_ progenies (238) was not consistent with the expected 3:1 wild-type:mutant phenotype segregation ratio of a single recessive gene (X^2^ = 52.71, *P* <0.0001), suggesting that multiple genes may be responsible for the Cadenza0900 phenotype. In contrast, the segregation pattern of the Cadenza0062 F_2_ population (51 plants) is consistent with the 3:1 mutant:wildtype segregation ratio expected of a single dominant gene (X^2^ = 0.53, *P* = 0.4669), suggesting that the Cadenza0062 phenotype is caused by a single dominant gene.

## DISCUSSION

### Use of sequenced mutant population to characterise RSA genes in wheat

Mutant analyses have played a key role in the identification of genes controlling key stages of root development. For instance, most of the genes identified to control root architecture in maize were identified via mutant analyses (Hochholdinger et al. 2018; Marcon et al. 2013). These include *RTCS, RTCL, RUM1* and *BIGE1* which display seminal root phenotypes (Taramino et al. 2007; Suzuki et al. 2015; von Behrens et al. 2011; Xu et al. 2015). Despite the buffering effect of genetic redundancy that often masks single homoeolog mutations in polyploid wheat (Borrill et al. 2015), our study highlights the usefulness of forward genetic screens to identify heritable variation for root development traits in wheat. These results are also consistent with recent examples of dominant mutations being identified in forward screens of the exome-sequenced populations (Harrington et al. 2019; Mo et al. 2018).

The use of a sequenced mutant population in this study also provided the opportunity to examine the presence of mutations in candidate genes from other species. For example, the phenotypes of the lower root number mutants (Cadenza0062, Cadenza0369, Cadenza0393, Cadenza0465, Cadenza0818, and Cadenza1273) are similar to the phenotypes of maize *rtcs* and *rtcl* mutations (Taramino et al. 2007), and their orthologous rice mutants (Liu et al. 2005; Inukai et al. 2005). *In-silico* examination of the coding regions in these mutants on the Ensembl Plant database revealed that Cadenza1273 contains a functional mutation in *TraesCS4D01G312800*, one of the three wheat homoeologs of *RTCS, RTCL* and *ARL1/CRL1*. Cadenza1273 harbours a G765A mutation in *TraesCS4D01G312800* producing a premature termination codon which results in a truncated 265 amino acid (aa) protein instead of the 289 aa native protein. Further molecular and genetic characterisation will be required to test if the G765A mutation in Cadenza1273 is responsible for its phenotype. However, this exemplifies the power of combining the sequenced mutant information with known candidate genes and a now fully annotated wheat genome (Appels et al. 2018). The other lower root number mutants do not contain any functional EMS mutations in the three wheat homoeologs of *RTCS, RTCL* and *ARL1/CRL1* and therefore likely represent new variation controlling seminal root development in cereals.

Similarly, the higher seminal root number phenotype of Cadenza0900 is similar to the phenotype of maize *bige1* (Suzuki et al. 2015) mutants. However, unlike *bige1* which is monogenic, we show that the phenotype of Cadenza0900 is most likely conditioned by more than one mutation. Also, *in-silico* examination of mutations in Cadenza0900 show that it does not harbour any mis-sense or non-sense mutations in the coding sequences of the three wheat orthologs of *BIGE1* (*TraesCS4A02G350200, TraesCS5B02G522900 and TraesCS5D02G521600*) thereby reducing the likelihood that the Cadenza0900 phenotype originates from mutations of the wheat *BIGE1* gene. It is, however, important to note that these *in-silico* investigations are restricted to mutations in the coding region of the wheat genome and we cannot rule out that mutations in promoter regions of these candidate genes might be responsible for some of the altered root number mutants identified.

### Relationship between grain size and seminal root traits

Size stratification of seeds in our study allowed an examination of the relationship between grain size and root architecture. We observed a positive effect of grain size on root length and number. Irrespective of the genetic background (wild-type Cadenza or mutant background), large-sized grain showed longer root length compared to medium-sized grain consistent with the rationale that bigger grains have greater carbohydrate reserves in their endosperm to support faster root elongation. We also noticed a weak but positive effect of grain size on the number of seminal roots developed in the mutant population. Grain size effect on seedling traits, including seminal root traits, can be attributed to constituent components – embryo and/or endosperm (Bremner et al. 1963; Meyer 1976). Interestingly, four of the lower root number mutants (Cadenza0393, Cadenza0465, Cadenza0818 and Cadenza1273) have smaller embryos compared to Cadenza. However, without further genetic analyses, it is premature to conclude that the small embryo of these mutants directly affects their seminal root number phenotypes. Indeed, grain size only accounts for a small proportion (3.4%) of the total variance in root number in our study, indicating that seed size *per se* is not a major determinant of root number. This is further underlined by the fact that differently sized wild-type Cadenza seeds show similar root number averages (Figure 3B).

Although informative, the qualitative stratification of grain size (large, medium and small) adopted in this study does not allow a quantitative modelling of grain size effect on root traits. We propose that a finer calibration and partitioning of grain size measurement into constituent parameters (width, length, height) and tissue (embryo and endosperm) components (Brinton and Uauy 2019) will allow for a finer understanding of the effects of these seed size components on root architecture.

### High-throughput platforms enables population-scale screening

The high-throughput afforded by the 2D platform used in Screen A in this study enabled the screening of a large mutant population (645 lines). Based on a previous 2D phenotyping pipeline implemented by Atkinson et al. (2015), our platform increases the throughput from 360 to 1,800 seedlings while maintaining a similar phenotyping rate (~4.5 minutes per plant). A similar high-throughput phenotyping platform was also recently adapted by (Adeleke et al. 2019) with a maximum phenotyping capacity of 672 plants. It is also worth noting that our high-throughput platform is relatively inexpensive and easy to implement using widely-available low-cost materials: storage boxes, germination paper, plastic sheets and paper binder clips. The entire phenotyping pipeline costed £5,570 to phenotype 6,240 plants: £3,280 to set-up the infrastructure and £2,090 recurring cost (paper and nutrient solution reagents). Besides the initial set-up cost, this represents a recurring cost of £0.33 per plant, making it cheaper than commercially available CYG germination papers. Also, we believe an important feature of this set-up is the modularity of phenotyping it offers, that is, the ability to set up experiment in small individual phenotyping units (boxes, Figure 2B and C) which can be used as experimental blocks. This modularity also allows flexibility of phenotyping scale as the experiments can be scaled up or down by adjusting the number of boxes used.

More than half of the processing time in Screen A was spent on image analysis. This included semi-automated analysis of RSA using RootNav as individual root tips need to be selected first before the root outline is automatically extracted. This processing time can be further improved by adopting the latest deep machine-learning algorithm recently applied to root image processing allowing fully automated extraction of root outline. This innovation will drastically reduce the time taken for image processing without compromising on accuracy (Pound et al. 2017).

Despite the high plasticity (e.g. dynamic response to varying water, nutrient and environmental conditions) associated with root traits, we obtained a high heritability estimate for seminal root number measurements across the two screens, similar to previous estimates (Maccaferri et al. 2016; Ma et al. 2017). While this high heritability may be due to the controlled hydroponic environment used in these experiments (Figure 2), it nonetheless demonstrates that seminal root number is a stable phenotype under strong genetic control and can be targeted for selection to improve RSA in wheat breeding programmes. There is also evidence to suggest that seminal root number phenotypes observed in hydroponic conditions are transferrable to soil conditions (Richard et al. 2015; Watt et al. 2013) and might therefore be useful under field conditions especially during the vegetative phase. We are currently evaluating the mutant lines under field conditions to determine this.

### Developmental and genetic characterisation of seminal root formation

Natural variation in wheat seminal root number has been shown to originate from defects in root primordia development in the embryo and/or differential activation of developed primordia to form fully emerged seminal roots (Golan et al. 2018). Our anatomical characterisation of intact non-imbibed seeds of the lower root number mutants highlight a tendency for these mutants to develop less than five root primordia, whereas wild-type Cadenza plants consistently develop five root primordia. In addition, these mutants form even fewer numbers of seminal roots than the root primordia they developed, suggesting that some of the primordia developed in these mutants might be dormant/inactive or arrested soon after activation. Together these results highlight both primordia development and activation as two important development check points in the formation of seminal root in wheat.

All the altered root number mutants develop SR_1_ and SR_2,3_ but show defects in the development of SR_4,5_ as in the lower root number mutants, or develop an extra root (SR_6_) as in Cadenza0900. This suggests that the development of the different seminal root types is under distinct genetic control. We did not recover mutant lines defective in either SR_1_ nor SR_2,3_: this could be due to the fact that defects in these root types might have severe negative or lethal effects on seedling growth. Golan et al. (2018) show that SR_1_ and SR_2,3_ contribute more to water uptake in durum wheat than SR_4,5_ under well-watered conditions. It is however possible that SR_4,5_ and indeed SR_6_ may contribute significantly to nutrient and water uptake under resource-limiting conditions where an increase in root surface area maximises soil exploration. Detailed field physiological evaluation will be required to better understand the cost-benefit relationship of the altered seminal root phenotypes.

The lower root number mutant Cadenza0062 shows a dominant mode of action with a 3:1 segregation ratio in the F_2_ population, characteristic of a monogenic trait. Until this and the other mutants are independently characterised, we cannot conclude that this dominant monogenic behaviour is representative of all the lower root number mutants, nor can we rule out the possibility of some of these mutations being allelic. Unlike Cadenza0062, the extra seminal root number mutant Cadenza0900 shows a recessive, multigenic phenotype that suggests a more complex genetic regulation of additional seminal roots in wheat. More detailed genetic characterisation and mapping will be required to better dissect the genetic control of these phenotypes.

### Outlook

Our work provides a complementary approach to the use of natural variation in dissecting the genetic control of seminal root development in wheat. The isolation of these mutants represents an important step in identifying the genetic determinants controlling wheat seminal root development and growth. These will be followed by extensive genetic characterisation to map these mutations to defined chromosomal positions and identify the causal gene(s) underlying their phenotypes. Given the high mutation rate in the Cadenza mutant population (~33 mutation per Mb; Krasileva et al. 2017), it will also be necessary to reduce the mutation load in these mutants to enable specific characterisation of each mutation. This can be achieved through backcrossing each mutant to wild-type Cadenza and selection of F_1_ progenies (for dominant mutations like in Cadenza0062) or F_2_ progenies (for recessive mutations like Cadenza0900) that retain the altered seminal root number phenotype (Uauy et al. 2017). These backcrossed mutants will also provide the background for developing crosses to examine interactions between the different seminal root number mutations.

## ACKNOWLEDGEMENTS

We would like to thank Dr Jonathan Atkinson, Prof Malcolm Bennett, Chloe Riviere and Guiditta Giordani for technical advice and assistance in the setting up the high-throughput 2D root screen and Andrew Davis for help with root imaging. This research is supported by the UK Biotechnology and Biological Sciences Research Council (BBSRC) Designing Future Wheat programme (BB/P016855/1), a Royal Society FLAIR award (FLR\R1\1918500) to OS, a BBSRC DTP award to RK, the Chief Scientist of the Israel Ministry of Agriculture and Rural Development grant (#12-01-0005), and the U.S. Agency for International Development Middle East Research and Cooperation grant (# M34-037).

