## Supplemental Figures for "Genetic screening for mutants with altered seminal root numbers in hexaploid wheat using a high-throughput root phenotyping platform"

### Slide 1
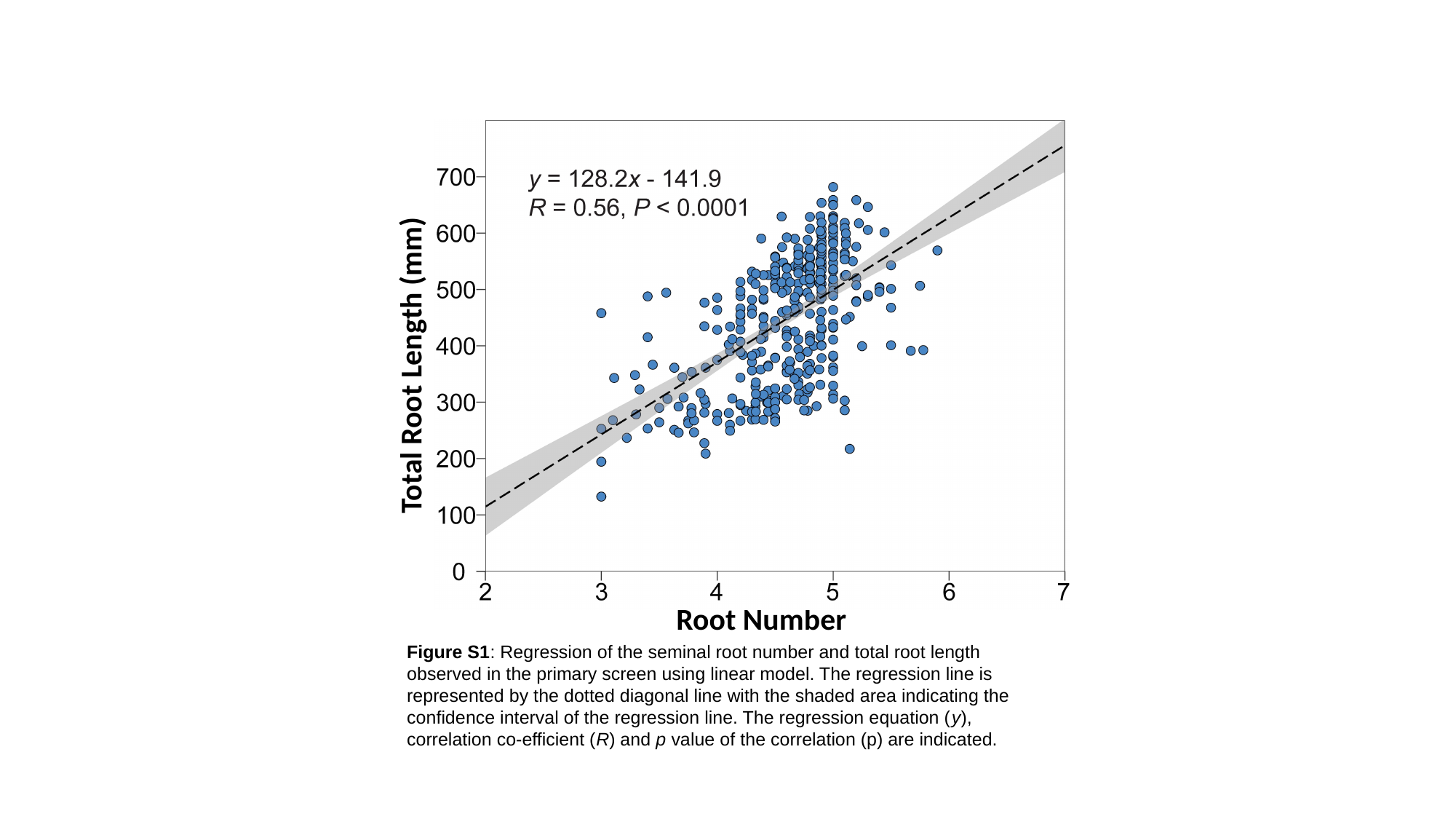

Total Root Length (mm)
Root Number
Figure S1: Regression of the seminal root number and total root length observed in the primary screen using linear model. The regression line is represented by the dotted diagonal line with the shaded area indicating the confidence interval of the regression line. The regression equation (y), correlation co-efficient (R) and p value of the correlation (p) are indicated.

### Slide 2
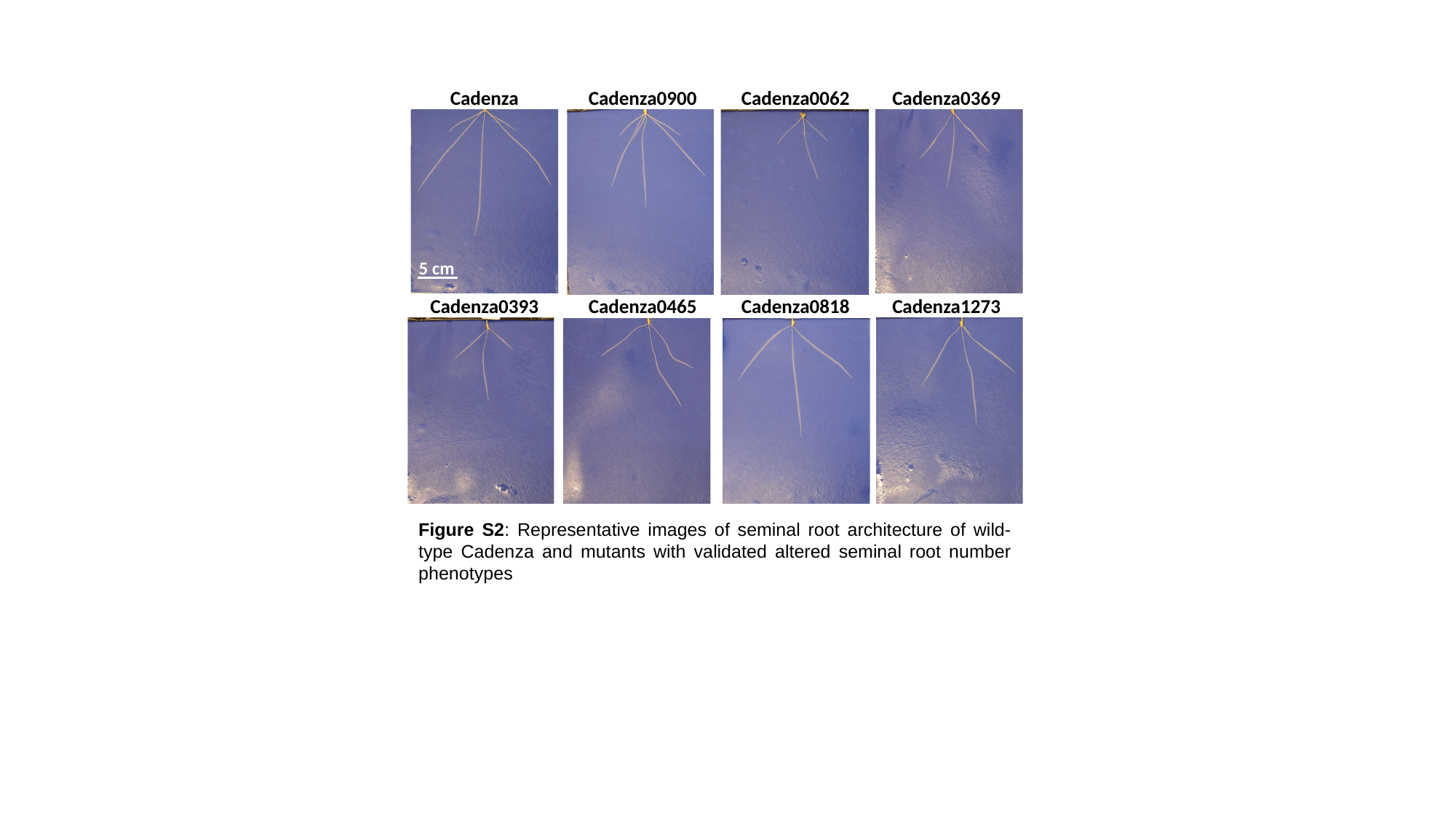

Cadenza
Cadenza0900
Cadenza0062
Cadenza0369
5 cm
Cadenza0393
Cadenza0465
Cadenza0818
Cadenza1273
Figure S2: Representative images of seminal root architecture of wild-type Cadenza and mutants with validated altered seminal root number phenotypes

### Slide 3
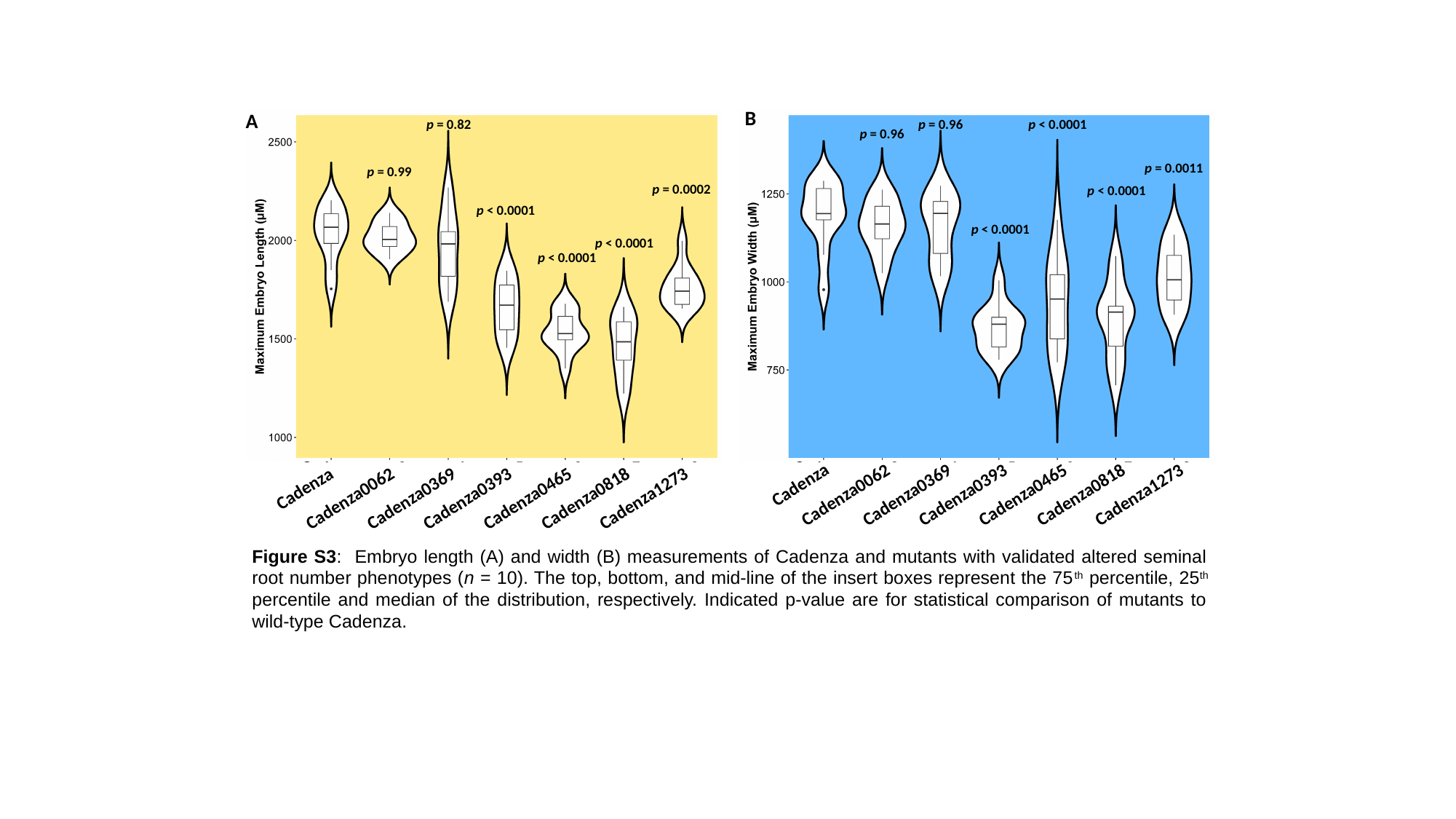

B
A
p = 0.96
p = 0.82
p < 0.0001
p = 0.96
p = 0.0011
p = 0.99
p = 0.0002
p < 0.0001
p < 0.0001
p < 0.0001
p < 0.0001
 p < 0.0001
Cadenza
Cadenza0062
Cadenza0369
Cadenza0393
Cadenza0465
Cadenza0818
Cadenza1273
Cadenza
Cadenza0062
Cadenza0369
Cadenza0393
Cadenza0465
Cadenza0818
Cadenza1273
Figure S3: Embryo length (A) and width (B) measurements of Cadenza and mutants with validated altered seminal root number phenotypes (n = 10). The top, bottom, and mid-line of the insert boxes represent the 75^th^ percentile, 25^th^ percentile and median of the distribution, respectively. Indicated p-value are for statistical comparison of mutants to wild-type Cadenza.
